# Biolistic transformation of *Haematococcus pluvialis* with constructs based on the flanking sequences of its endogenous alpha tubulin gene

**DOI:** 10.1101/420711

**Authors:** Guanhua Yuan, Wenlei Zhang, Xiaoying Xu, Wei Zhang, Yulin Cui, Tianzhong Liu

**Author notes:** Correspondence author: **E-mail** (WZ); (TL).

## Abstract

The complete sequence information of the alpha tubulin (*tub*) genes was obtained from both *Haematococcus pluvialis* NIES144 and SCCAP K0084., Putative transcriptional elements and polyadenylation signals were identified respectively in their 5’ and 3’ flanking regions. Three selection cassettes of tub/aadA, tub/hyr and tub/ble with 3 different antibiotic-resistant genes fused between the 5’ and 3’ flanking sequences of the *tub* gene were constructed and utilized for biolistic transformation of *H.pluvialis*. Antibiotic resistant transformants were obtained in the bombardments with the tub/aadA cassette in 2 strains. It was found that, the foreign tub/aadA DNA could be completely transferred and inherited in their genome through non-homologous recombination. Moreover, transcripts of the insert and spectinomycin resistance were identified. Transformation efficiencies up to 3×10^-5^ per μg DNA could be obtained in *H.pluvialis* NIES144 or SCCAP K0084 through utilization of a culture with a high percentage of flagellate cells and by optimizing bombarding protocol. The presented selection marker and optimized transforming procedures in this report should strengthen the platform technology for genetical manipulation and modification of *H.pluvialis*.

## Introduction

Astaxanthin (3, 3’-dihydroxy-ß-carotene-4, 4’-dione) is the strongest antioxidant in the nature, and has been widely used as an additive in health care products and cosmetics [1]. *Haematococcus pluvialis* has attracted great attention as its richest biological source. Currently, *H.pluvialis* cultivation has been commercialized to produce astaxanthin powder, which can be utilized as feedstocks or as additives in aquacultural, pharmaceutical and nutraceutical products [2]. *H.pluvialis* has been considered as an organism for the production of recombinant pharmaceutical proteins[3].

Researches on *H.pluvialis* have taken place over several decades, mostly focused on cell differentiation [4], astaxanthin metabolism[5], cultivation methods [6] and astaxanthin extraction technology[7]. Continuous efforts were made to improve its cellular growth and the level of astaxanthin accumulation through traditional mutagenesis of *H.pluvialis* [8]; however there are no reports of phenotypic modification through genetic engineering methods. To date, only 4 researchers have reported stable genetic transformation of *H.pluvialis*. Steinbrenner & Gerhard Sandmann in 2006 [9] transformed *H.pluvialis* with a mutated phytoene desaturase (*pds*) gene conferring resistance to norflurazon, using a biolistic method. Sharon-Gojman et al. in 2015[10] optimized this vector system by shortening the promoter sequence from 2000 base pairs (bp) to 1000 bp. And an agrobacterium-mediated method using a plant-general plasmid pCAMBIA1301 could transform *H.pluvialis*, with a foreign *hptII* gene been identified in the transformants [11]. Successful chloroplast transformation was also achieved through biolistic bombardment of *H.pluvialis* using a plasmid pHuplus, which possessed the *aadA*, an aminoglycoside-3’-adenylyltransferase that provides resistance to streptomycin and spectinomycin, driven by an endogenous large subunit of ribulose bisphosphate carboxylase (rbcL) promoter and terminator [12]. However, compared with the model algae species *Chlamydomonas reinhardii*, reports of high-efficient genetic transforming and expression systems are still scarce, which has limited their application to the directed genetic modification in *H.pluvialis*.

In order to construct more efficient screening markers for transformation of *H.pluvialis*, the endogenous 5’ and 3’ flanking sequences of the alpha tubulin gene were cloned and 3 antibiotic-resistant genes were each independently fused between them. The resulting fusion cassettes were used to biolistically transform of *H.pluvialis*. The transforming efficiency, the influencing factors, the integration and expression of the introduced DNA fragment were investigated and discussed.

## Materials and Methods

### Strains and cultivation

*H.pluvialis* NIES144 and SCCAP K0084 were purchased from the Microbial Culture Collection at the National Institute for Environmental Studies and the Scandinavian Culture Collection of Algae and Protozoa, respectively. Both strains were maintained in BG11 medium at 25 °C with continuous illumination of about 25-30 μmol m^-2^ s^-1^.

### Basic Polymerase chain reaction (PCR) cloning and analysis

The regular PCR was performed as follows: a 10 min initial denaturation step at 95 °C, followed by 30 cycles of denaturation for 30s at 95 °C, annealing 30s at 57 °C, extension 1min per kilo bp sequence length at 72 °C, and a final 10 min extension step at 72 °C. The TransTaq DNA polymerase High Fidelity (HIFI) kit (TransGene Biotech, Beijing, China) and 2× PCR BestaqTM MasterMix (Applied Biological Materials Inc. Vancouver, Canada) were utilized for DNA cloning and PCR analysis, respectively. The sequence information of the primers was provided in Table 1. The pEASY-T1 Cloning Kit (TransGene Biotech, Beijing, China) was utilized for TA cloning of the target sequence.

**Table 1.**
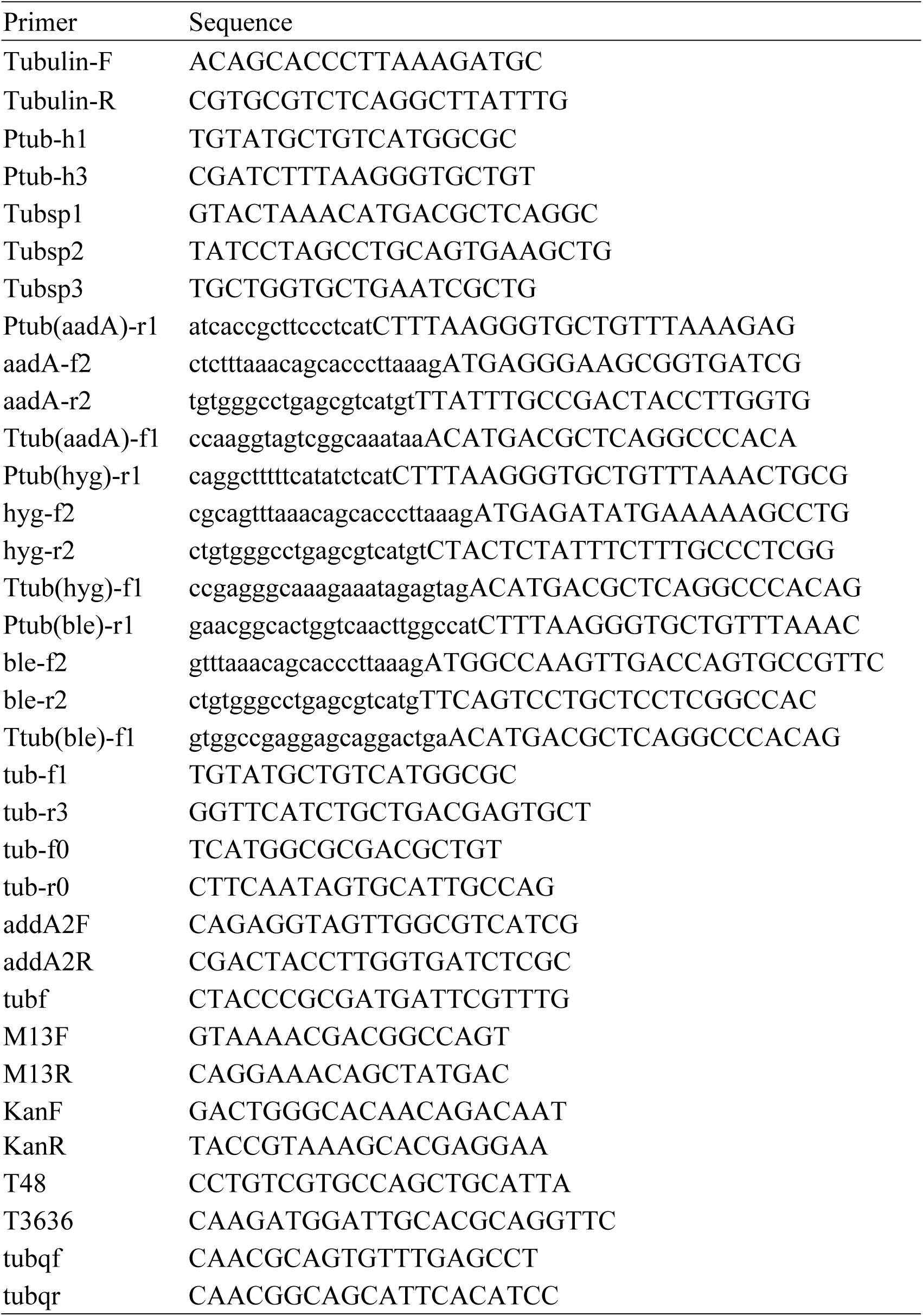
The sequence information of the primers

### Isolation of genomic DNA

Genomic DNA of *H.pluvialis* was isolated using the PlantZol reagent (TransGene Biotech, Beijing, China). The harvested *H.pluvialis* cells were firstly mixed with the PlantZol reagent. Then, the suspension was grounded with an electric grinder (G506001, Sangon Biotech, Shanghai) on ice and incubated at 55 °C for 15 min. After extraction with 1 vol of isopyknic phenol:chloroform:isoamyl alcohol (25:24:1), the genomic DNA was precipitated from the supernatant with 1 vol isopropyl alcohol. The genomic DNA precipitate was dissolved with distilled water at 55 °C after washing with 70% alcohol and was stored in −20 °C for next use.

### Extraction of total RNA and cDNA synthesis

Total RNA of *H.pluvialis* was extracted using a RNAiso Plus kit (TaKaRa Bio Inc., Kusatsu, Japan). Harvested *H. pluvialis* paste, previously frozen with liquid nitrogen, was ground in a mortal and then mixed with RNAiso Plus reagent. The homogenate was transferred into a 1.5 mL centrifuge tube and incubated for about 5 min at room temperature. After an extraction for 5 min with 1/5 vol chloroform, the RNA was precipitated with isopropyl alcohol. The RNA precipitate was dissolved with RNase-free distilled water after washing with 80% alcohol and stored at −70 °C. First-stand cDNAs was synthesized using a PrimeScript TM II 1st Strand cDNA Synthesis Kit (TaKaRa Bio Inc., Kusatsu, Japan). RNA sample was firstly mixed with the Oligo dT primer and dNTP mixture in a microtube. The mixture was kept at 65 °C for a 5 min denaturing reaction and then placed on ice to cool. Then, RNase-free dH_2_O, 5× PrimeScript II Buffer, RNase inhibitor and PrimeScript II RTase were added to the mixture. After a two-step reaction of 60min at 42 °C and 15min at 70 °C, the resulting cDNA sample was stored at −70 °C.

### Cloning of the alpha tubulin (*tub*) gene and its flanking sequences

PCRs were performed with the primer pairs of tubulin-F/tubulin-R and Ptub-h1/Ptub-h3 to obtain the *tub* genes and their 5’ flanking sequences in the 2 *H.pluvialis* strains of NIES144 and SCCAP K0084. Both the genomic DNA and cDNA were used as template. These primers were designed based on the sequence data from accession numbers of AY894136.1 and GQ470501.1 of Genebank. The PCR products were cloned into pEASY-T1 vectors and sequenced at the Shanghai Personal Biotechnology Corporation after transformation into *Escherichia coli* DH5α.

In order to obtain the 3’ flanking sequences of the alpha-tubulin without any available information on homologous sequences, a partially overlapping primer-based PCR was used for genome walking [13], with a Genome Walking Kit (TaKaRa Bio Inc., Kusatsu, Japan). Three specific synthetic primers of Tubsp1, Tubsp2 and Tubsp3 were designed. Each primer had a relatively high annealing temperature. Three rounds of PCR were performed for each walking process using the product of the previous PCR as a template for the next PCR. Each PCR reaction mixture had 1×LA PCR Buffer II(Mg^2+^ plus) containing 0.4mM dNTP mixture, 2.5 U TaKaRa LA Taq, 0.2 μM of each primer and a certain amount of template. The first round of PCR had 3 anneling stages: stage 1 was 5 high-stringency (65 °C) cycles; stage 2 was 1 low-stringency (25 °C) cycle; stage 3 was 30 high-stringency (65 °C) cycles. And the next 2 rounds of PCR had 2 annealing stages: stage 1 was 30 high-stringency (65 °C) cycles; stage 2 was 15 low-stringency (44 °C) cycles. After agarose gel electrophoresis and gel extraction, the fragments were cloned into pEASY-T1 vectors and sequenced.

### Construction of transforming vector

A fusion PCR method [14] was performed to construct the cassettes of tub/aadA, tub/hpt and tub/ble, in which 3 antibiotic-resistant genes, *aadA, hptII* and *sh ble* were independently fused with the *tub* promoter and terminator of *H.pluvialis* NIES144 as shown in Fig. 1.B. The *aadA* was obtained from the plasmid pHuplus [12], the *hptII* from the plasmid pCAMBIA1305 (Cambia, Australia) and the *sh ble* gene from the plasmid pPha-T1 [15]; these genes conferred resistance to spectinomycin, hygromycin and zeomycin. Taking the construction of the tub/aadA cassette as an example, 3 target fragments of Ptub, aadA and Ttub were obtained by regular PCR with the primer pairs of tub-f1/Ptub(aadA)-r1, aadA-f2/aadA-r2 and Ttub(aadA)-f1/tubr3, respectively. The products were purified by agarose gel electrophoresis using a Agarose Gel DNA Extraction Kit (TaKaRa Bio Inc., Kusatsu, Japan). The 3 purified DNA fragements in equal amounts (200-800 ng) were mixed with pfu DNA polymerase and dNTPs in a 50 μL PCR reaction system. This mixture underwent with a PCR fusion procedure as follows: a 10 min initial denaturation step at 95 °C, followed by 13 cycles of denaturation for 30s at 95 °C, annealing 30s at °C, extension 1min per kilo bp sequence length at 72 °C, and a final 10 min extension step at °C. The final mixture was utilized as a template for PCR amplification of the target tub/aadA fragment with the primer pair of tub-f0/tub-r0. The resulting fusion fragments were cloned into the plasmid pEASY-T1 by TA cloning. The sequence of the fusion fragments was verified by plasmid sequencing. Three plasmids were named as pEASY-tub/aadA, pEASY-tub/hyr and pEASY-tub/ble, based on their respective antibiotic resistance gene fusion. Their sequences were submitted to Genebank with the accession numbers of MH752993, MH752994 and MH752992, respectively.

**Fig 1.**
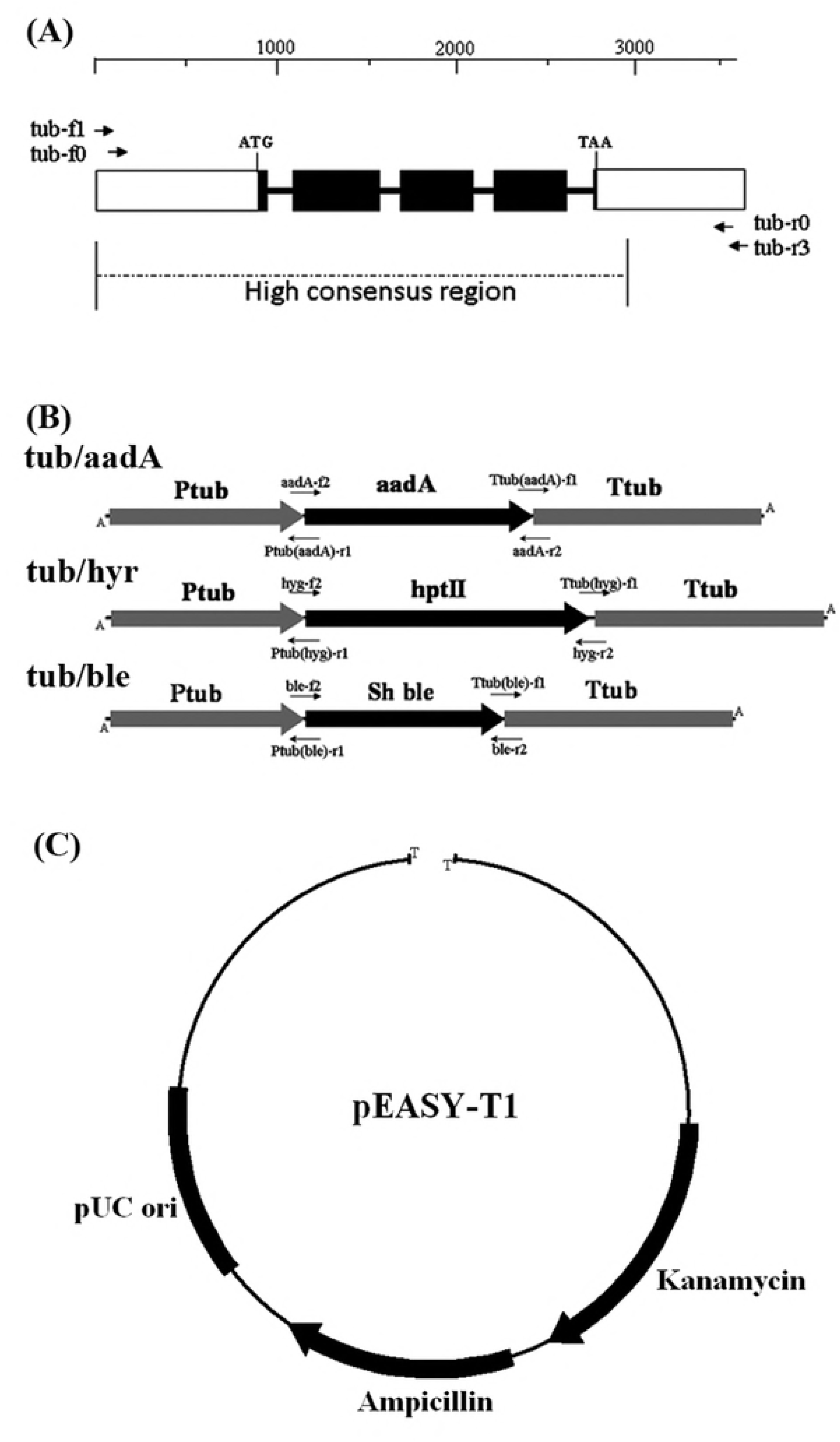
Maps of the alpha tubulin gene from the geonome of *Haematococcus pluvialis* (A), the fusion cassettes of tub/aadA, tub/hyr and tub/ble (B), and the T clone vector of pEASY-T1 (C). The dark boxes, thicken lines and blank boxes in (A) represented the exons, introns and 5’ or 3’ flanking regions of the alpha tubulin gene. The ruler of nucleotide numbers and dash line above and below them indicate the sequence lengths and high sequence consensus, respectively. The dark arrows showed the sites of the primers utilized in construction of the 3 fusion cassettes.

### Transformation of *H.pluvialis*

For biolistic transformation, *H.pluvialis* cells were grown in BYA medium as described previously [16]. Cultivation times were chosen based on their growth and the percentage of flagellate cells. Cells were harvested by centrifugation and suspended in BG11 medium to a cell density of 2-3×10^8^ ml^-1^. A 100 μl aliquot of cell suspension was layered onto a 25mm mix cellulose ester filter membrane, which was placed on a BG11 solid plate for bombardment. The bombardment was performed with a PDS-1000/He Biolistic Particle Delivery system (Bio-Rad, USA) as described previously [9]. Plates were bombarded from a distance of 6 cm using 1350 psi rupture disks. DNA coated particles were prepared by mixing of 50 μl M 17 tungsten particle solution (60 mg/ml in H_2_O) with 5 μl of a DNA solution (>1 μg/μl), 50 μl of 2.5 M CaCl_2_, and 20 μl of 0.1 M spermidine base. This was followed by a 10min incubation on ice and 2 centrifugal washes with 70% and then 100% ethanol before final resuspension in about 50 μl ethanol. Ten μl of prepared DNA-coated particle solution was layered on a macrocarrier for 1 bombardment. After the bombardment, the cells were washed from the filter membrane into about 25 ml of BG11 liquid medium for the incubation at 25 °C with continuous illumination at 10-15 μmol m^-2^ s^-1^. After 48 hr regeneration, *H.pluvialis* cells were screened on solid plates of antibiotic-containing TAP media at 25 °C with continuous illumination of about 25-30 μmol m^-2^ s^-1^. For transformation with tub/aadA, tub/hyr and tub/ble, 200 μg ml^-1^ of spectinomycin, 10 μg ml^-1^ of hygromycin and 5 μg ml^-1^ zeocin were added to the respective media.

### Determination of cell density, flagellate percentage and viability

The cell density, flagellate percentage and viability were determined by hemocytometer counting using an Olympus BX53 microscope. The number of flagellate cells was judged visually by its flagella and cell morphology. Phenosafranine staining was used to determine cell viability; cells that stained into red were counted as dead. All results were obtained from ≥ 200 cells in repeated samples.

### Sequence analysis

Sequence alignments were performed with Clustal X. The putative cis-regulatory elements of the promoter were analyzed using the PlantCARE databases (http://bioinformatics.psb.ugent.be/webtools/plantcare/html/). The nucleotide sequences of the tubulin genes along with their flanking regions from *H.pluvialis* NIES 144 and SCCAP K0084 were deposited in the GenBank database under accession numbers MH752990 and MH752991.

## Results

### Similarity of the alpha tubulin (*tub*) gene and its flanking sequence in 2 *H.pluvialis* strains

The *tub* and its flanking sequences were respectively cloned and sequenced from the genomes of *H.pluvialis* NIES144 and SCCAP K0084. Their DNA sequences, with lengths of 3663 bp and 3430 bp, were obtained. Fig 1.A shows their features. The *tub* gene sequences, consisted of 5 exons and 4 introns, along with the whole 5’ flanking region and part of the 3’ flanking region, and there was a high level of sequence consistence between the 2 strains. The encoding region had a length of 1356 nts with differences between the 2 strains at several nucleotide sites. All 4 introns followed the GT-AG rule with 5’-GT and 3’-AG at their borders, with the same splicing sites for both strains. Respectively, 909 bp and 908 bp of 5’ flanking sequence was obtained for *H.pluvialis* NIES144 and SCCAP K0084, which were identical except for 2 divergent nucleotide sites. Based on the genomic walking strategy, 846 bp and 610 bp of 3’ flanking sequence was obtained for the strains, respectivley. Except for about 288 bp sequence after the termination codon, there was little or no in the rest of the 3’ flanking regions.

### Transcriptional elements in the 5’ flanking sequence of the *tub* gene

To identify the regulatory elements of the promoter, the 5’ flanking sequence of the *tub* gene was searched using the PLACE databases. One putative TATA-box was detected at −78bp from the ATG start codon and 11 putative CAAT-boxes were scattered from - 836bp to −134 bp, though none of them completely obeyed the consensus of GGCTCAATCT (Table 2). Other possible regulatory elements involved with light responsiveness were found, including G-box, ACE, CATT, ATCT, GAG and T-box motifs. These might function in the light-induced expression of the gene. There were also several other possible elements or motifs involved with specific expression, such as 3 meristem-specific elements including CAT-box and CCGTCC-box, 2 endosperm-specific skn-1 motif, 1 circadian-controlled element, 1 drought-induced MYB binding site(MBS) and 1 abscisic-responsive element(ABRE). All of them were conserved in 2 the *H.pluvialis* strains, suggesting they may be involved with gene expression related to environmental or stress responses.

**Table 2.** Transcriptional elements in the 5’ flanking region of the alpha tubulin gene from *H. pluvialis* NIES144 and SCCAP K0084.

| Element | Function | Count |
| --- | --- | --- |
| TATA-box | core promoter element around -30 of transcription start | 1 |
| CAAT-box | common cis-acting element in promoter and enhancer regions | 11 |
| A-box | cis-acting regulatory element | 1 |
| G-box | cis-acting regulatory element involved in light responsiveness | 4 |
| ACE | cis-acting element involved in light responsiveness | 2 |
| CATT-motif | part of a light responsive element | 1 |
| ATCT-motif | part of a conserved DNA module involved in light responsiveness | 1 |
| GAG-motif | part of a light responsive element | 1 |
| T-box | part of a light responsive element | 1 |
| CAT-box | cis-acting regulatory element related to meristem expression | 2 |
| CCGTCC-box | cis-acting regulatory element related to meristem specific activation | 1 |
| Skn-1_motif | cis-acting regulatory element required for endosperm expression | 2 |
| circadian | cis-acting regulatory element involved in circadian control | 1 |
| ABRE | cis-acting element involved in the abscisic acid responsiveness | 1 |
| MBS | MYB binding site involved in drought-inducibility | 1 |

### Elements potentially involved with mRNA 3’ end formation

For *H.pluvialis* NIES144 and SCCAP K0084, 846bp and 610bp DNA sequences were cloned at 3’ of the TAA termiantion codon, respectively. A conservative arrangement of polyadenylation signals similar to those found in plants was identified. A possible far-upstream element (FUE) with a TTGTAA sequence was located 175bp downstream of the TAA termination codon in both strains (Fig 2). In addition, at 303bp and 193bp downstream of the TTGTAA motifs, a possible near-upstream element (NUE), characterized by a typical ployadenylation signal with a consensus sequence of AATAAA, was found. At 10-30bp downstream of the NUEs, possible mRNA cleavage sites (CS) with the consensus sequence of CA could also be found. So the cloned 3’ flanking sequences might contain all the cis elements required for mRNA 3’ end formation.

**Fig 2.**
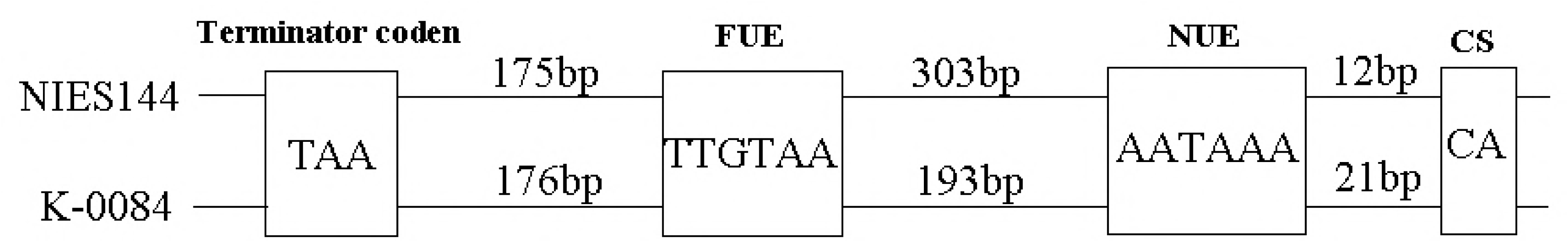
Plant-like polyadenylation signals in the 3’ flanking regions of *H.pluvialis* alpha tubulin genes. FUE, the Far-Upstream Element; NUE, the Near-Upstream Element; CS, the Cleavage/ polyadenylation Site

### Biolistic transformation of *H. pluvialis*

As shown in Fig 1.B, the *aadA, hptII* and *sh ble* cassettes, which encoded the spectinomycin, hygromycin and zeocin resistant genes, were respectively fused with 5’(Ptub) and 3’(Ttub) flanking sequences of the tubulin gene from *H.pluvialis* NIES144, using fusion PCR technology. The 3’ fusion fragments were then individually cloned into pEASY-T1 (Fig 1.C). The constructed plasmids pEASY-tub/aadA, pEASY-tub/hyr and pEASY-tub/ble were used for bombardments of *H.pluvialis* NIES144 and K0084. In every experiment, cells were bombarded with particles containing no DNA as a control sample. After bombardment and a subsequent 48 hours regeneration, the cells transformed with pEASY-tub/aadA, pEASY-tub/hyr and pEASY-tub/ble were screened for 3-4 weeks on the TAP plates with the addition of spectinomycin, hygromycin and zeocin, respectively. The growth of all of the control samples of H. *pluvialis* was totally inhibited on the TAP plates containing antibiotics. Colonies resistant to the corresponding antibiotic were obtained in repeated experiments of *H.pluvialis* NIES144 or SCCAP K0084 bombarded with the plasmid pEASY-tub/aadA or with the tub/aadA PCR fragment. No resistant colonies were obtained on attempted biolistic transformation with cassettes of either tub/hyr or tub/ble. Spectinomycin resistant clones were inoculated into fresh media for amplification PCR on extracted genomic DNAs with the *aadA*- specific primer pair of aadA2f/aadA2r. Fig 3 demonstrated that the foreign *aadA* gene could be detected in most of the clones obtained in different experiments.

**Fig 3.**
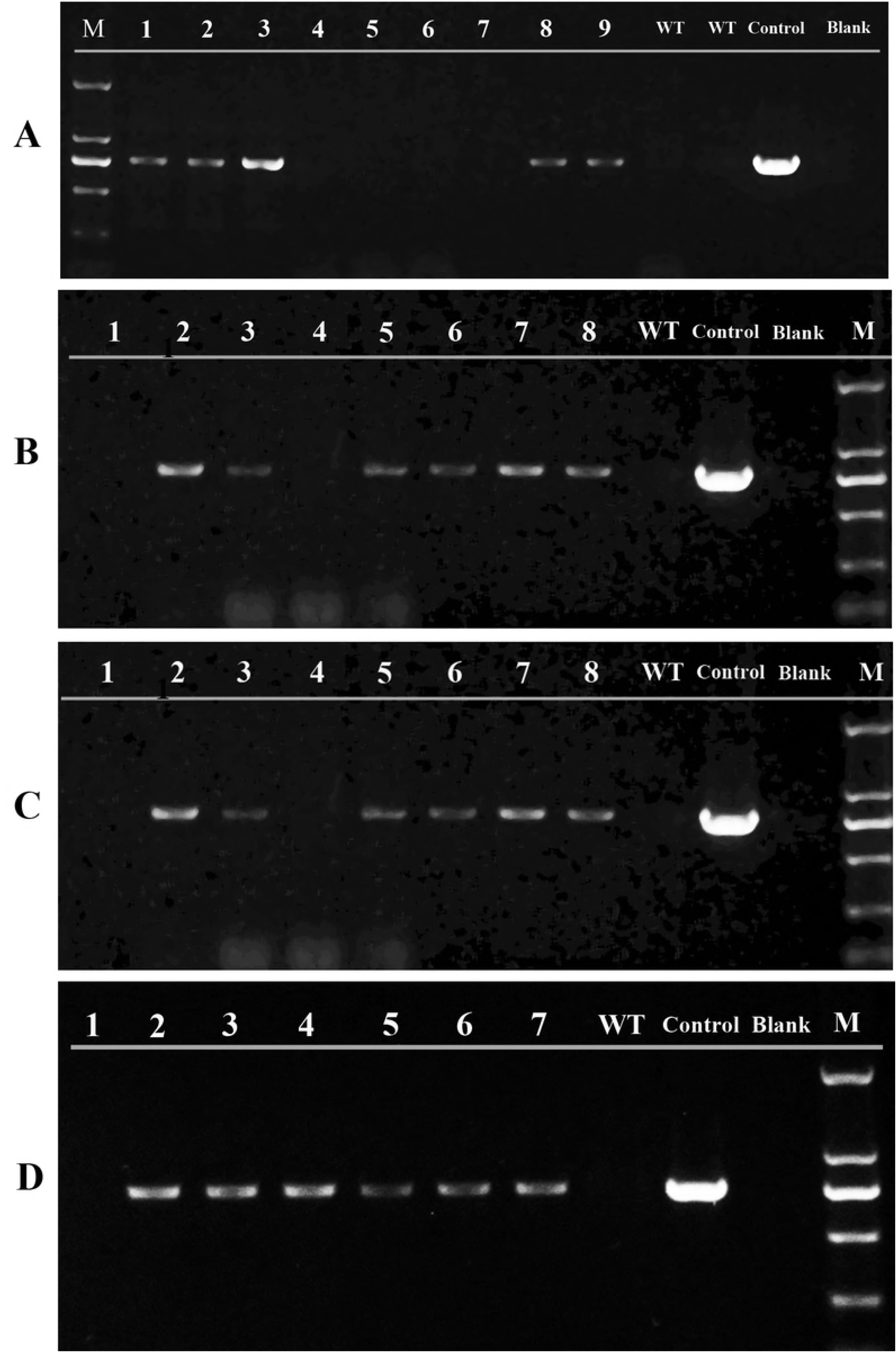
PCR detection of the *aadA* gene in the clones resistent to spectinomycin following separate bombardments of (A, C) *H. pluvialis* NIES 144 and (B, D) SCCAP K0084 with (A, B) pEASY-tub/aadA and the (C, D) the tub/aadA PCR fragment. The PCRs were performed with the primer pair aadA2f/ aadA2r using extracted genomic DNAs as a template. Lane M, DL 2000 DNA markers; Lane 1-9, independent spectinomycin resistant clones; Lane WT, negative control of wild type *H.pluvialis*; Lane Control, positive control of the plasmid pEASY-tub/aadA; Lane Blank, negative control of ddH_2_O

### Transforming efficiency and key influencing factors

Before biolistic transformation, *H.pluvialis* NIES144 and SCCAP K0084 were cultivated in BYA media. Their cellular growths and differentiation were recorded (Fig 4). With the extended cultivation days, both strains tended to convert from motile flagellates into nonmotile cells. *H.pluvialis* SCCAP K0084 showed more quicker growth and a higher converting tendency than strain NIES144. In order to avoid the influence of cell differentiation on transforming efficiency, logarithmic cultures after 6 days and 5 days cultivation of *H.pluvialis* NIES144 and SCCAP K0084, respectively, were chosen for biolistic bombardments. These logarithmic cultures had similar cell density of > 3×10^5^ ml^-1^ and a similar content of flagellate cells (> 80%). For each bombardment, about 3× 10^7^ cells was uniformly pipetted onto a cellulose filter membrane. After the bombardment, the cells needed to be immediately washed from the membrane and inoculated into fresh liquid media for a regenerating incubation, as long time incubation on the membrane led to high mortality (Fig 5) and ultimately failed recovery of transformants. After the regeneration, per aliquot with about 2×10^5^ cells was spreaded on 1 screening plate with a 6 cm diameter for about 3-4 weeks cultivation. Table 3 shows the yields of resistant colonies from different bombardments. At regular bombardment condition with about 1.18 μg DNA bombarded onto 3×10^7^ cells, 58.75 ± 13.39 and 49.86 ± 37.42 resistant colonies in average were obtained on the plates of pEASY-tub/aadA-transformed *H.pluvialis* NIES144 and SCCAP K0084, respectively. The total colony yields were 470 and 430, respectively, with a transforming frequency of 1.3×10^-5^ and 1.2×10^-5^ per μg DNA. When DNA amount was increased to about 1.7 μg per 3×10^7^ cells, average and total resistant colony yields elevated to 193.13±22.43 and 181.00 ± 40.81, 1545 and 1405, respectively. This corresponded to the higher transforming frequencies of 3.0×10^-5^ and 2.7×10^-5^ per μg DNA, respectively. The PCR fragment of the tub/aadA was also used for bolistic transformation instead of the plasmid DNA, however, lower transforming efficiencies of 0.4×10^-5^ and 0.9×10^-5^ per μg DNA were obtained for strains NIES144 and SCCAP K0084, respectively.

**Table 2.** Frequencies of resistant clonies achieved from different bombardments with the tub/aadA cassette

| DNA | Strain | DNA quantity($\mu\text{g}$)<sup>a</sup> | Average number of resistant clones<sup>b</sup> | Total number of resistant clones<sup>b</sup> | Clony frequency per $\mu\text{g}$ DNA<sup>c</sup> |
| --- | --- | --- | --- | --- | --- |
| pEASY-tub/aadA | NIES 144 | 1.71 | $193.13 \pm 22.43$ | 1545 | $3.0 \times 10^{-5}$ |
| | | 1.18 | $58.75 \pm 13.39$ | 470 | $1.3 \times 10^{-5}$ |
| | K0084 | 1.71 | $181.00 \pm 40.81$ | 1405 | $2.7 \times 10^{-5}$ |
| | | 1.18 | 49.86±37.42 | 431 | $1.2 \times 10^{-5}$ |
| Tub/aadA<br>fragment | NIES 144 | 1.03 | 19.83±9.75 | 119 | $0.4 \times 10^{-5}$ |
| | K0084 | 1.03 | 37.72±22.52 | 285 | $0.9 \times 10^{-5}$ |

a. Average DNA usage calculated from DNA concentration utilized in preparation of DNA-coated
particles
b. About $3 \times 10^7$ cells were bombarded for each experiment and screened on multiple solid plates
c. Total resistant clony numbers divided by cell numbers and DNA usage in one bombardment

**Fig 5.**
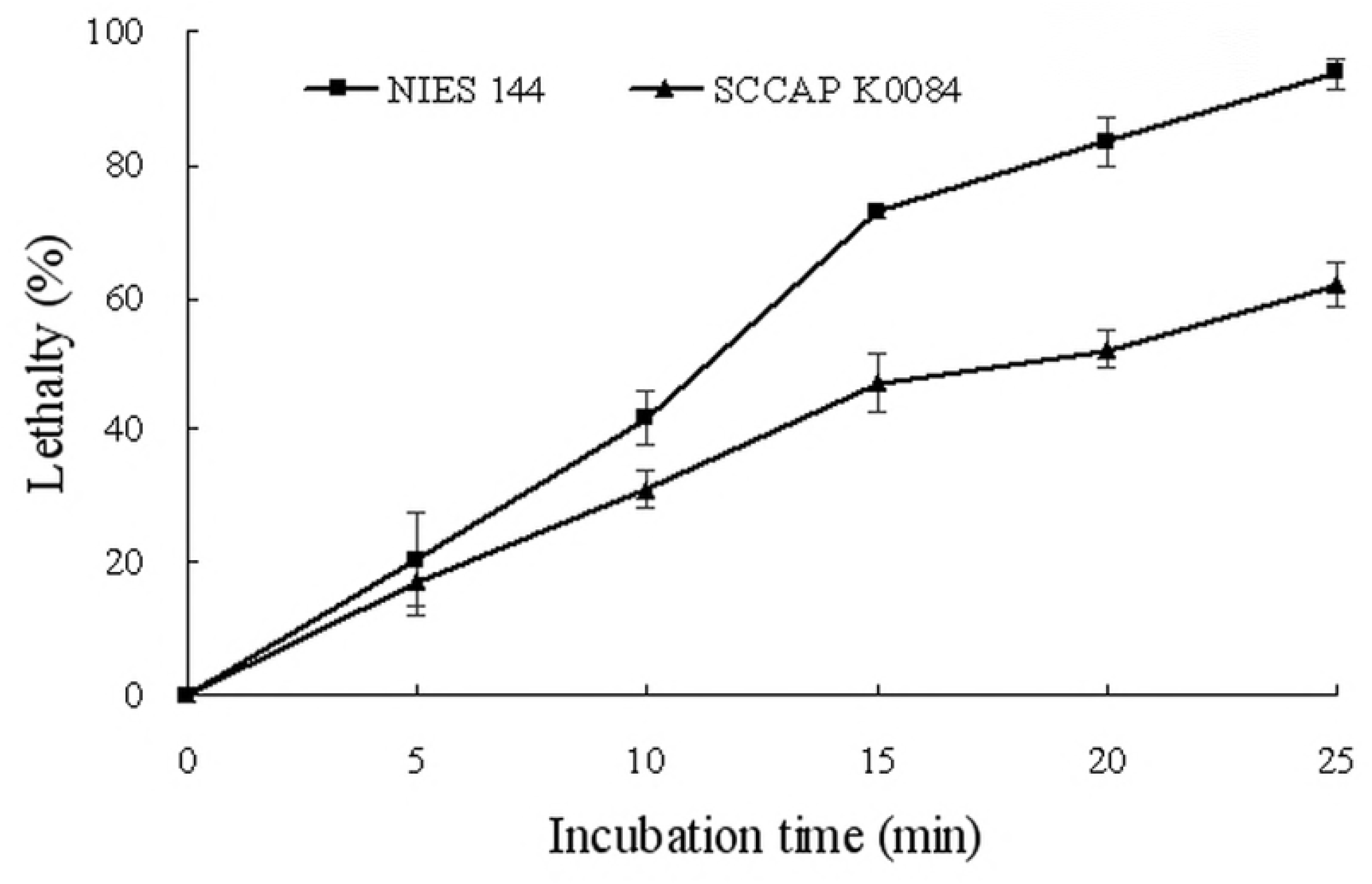
Relative lethality of concentrated *H.pluvialis* cells after prolonged incubation on membrane filter. All the data were from duplicate samples.

**Fig 4.**
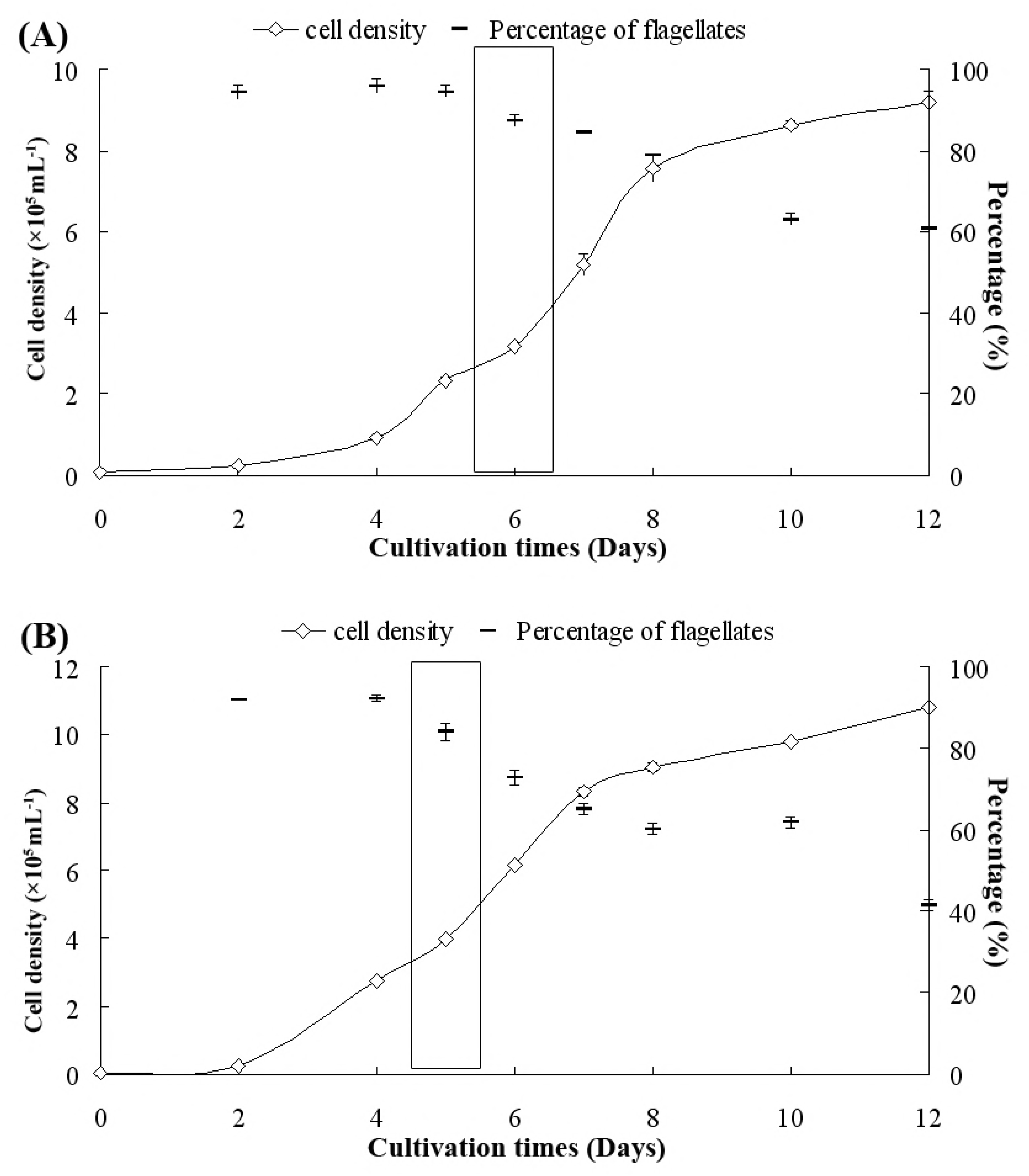
Growth and percentage of flagellate cells in the cultures of *Haemaococcus pluvialis* NIES144 (A) and SCCAP K0084 (B). The rectangular panel on each figure indicates the cultivation time chosen for preparation of *H.pluvialis* cells before bombardments. All the data were from duplicate samples.

### Integration of the foreign DNA into the *H.pluvialis* genome

To investigate how the foreign *aadA* gene was integrated into the *H.pluvialis* genome, the transgenic clones were examined by PCR with different primer pairs. Fig 6. A shows the binding sites of the primers on the *H.pluvialis* genome or on pEASY-tub/aadA. By analzing 5 independent transgenic clones from *H.pluvialis* NIES144 with the primer pairs, aadA2f/aadA2r, Tub-f0/aadA2r and aadA2f/tub-r0, it was demonstrated that the aadA gene drived by Ptub and Ttub sequences was completely integrated into the *H.pluvialis* genome (Fig 6.B). In addition, using the primer pair of tub-f0/tub-r0, the wild type tubulin gene and tub/aadA cassette was shown to simultaneously exist in the genomes of the 5 transgenic clones. Hmologous double crossover was not observed through the PCR from the primer pair of Tub-F1/Tub-R3, which located in the lateral sequences of Ptub and Ttub in the *H.pluvialis* genome. To identify possible single crossover events, 2 primer pairs of aadA2f/tub-R3 and tub-f1/aadA2r were utilized; however no gel banding indicative of single homologous crossover was observed in the transformants. To confirm the stablility of the transgenic colony, these transformants were sustained in media with and without spectromycin for at least 3 rounds of cultivation. For both strains, the tub/aadA cassette was stably maintained through generations.

**Fig 6.**
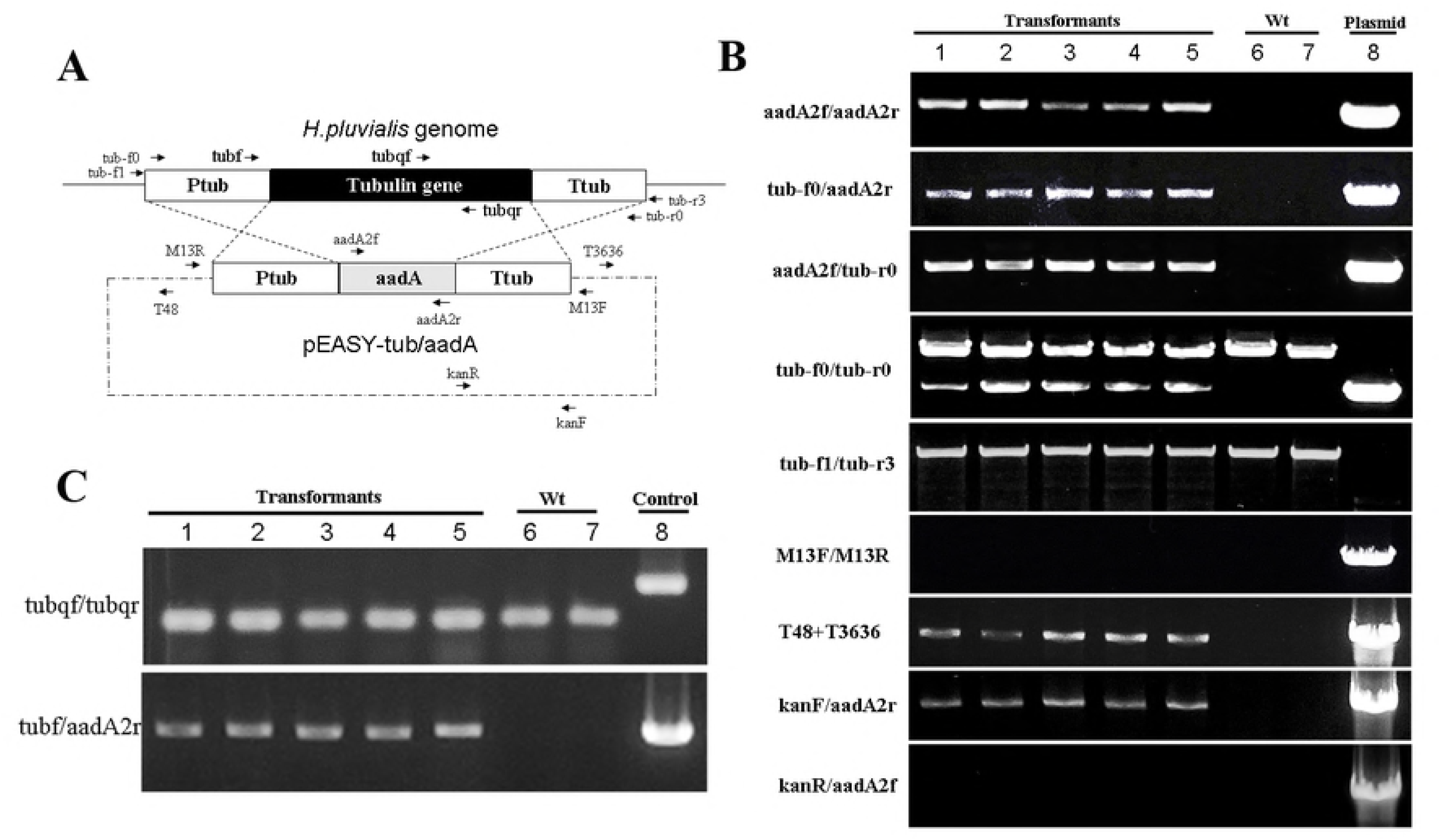
PCR analysis of genome integration and transcript expression of the pEASY-tub/aadA DNA in 5 independent transformants of *H.pluvialis* NIES144. (A) Map showing the homologous regions between *H.pluvialis* genome and the plasmid pEASY-tub/aadA (crossed dash lines), and the sites of primers used in the PCR analysis (dark arrows); (B) PCR identification of genome integration with different primer pairs in 5 transformants (Lanes 1-5), wild type *H.pluvialis* (Lanes 6-7) and the plasmid pEASY-tub/aadA (Lane 8); (C) PCR identification of transcript expression with different primer pairs in 5 transformants (Lanes 1-5), wild type *H.pluvialis* (Lanes 6-7) and the control samples of *H.pluvialis* genomic DNA and the plasmid pEASY-tub/aadA DNA for primer pairs of tubqf/tubqr and tubf/aadA2r, respectively (Lane 8).

### The expression of the *aadA* gene in the transformants

Transcription expression of the foreign *aadA* gene were identified in 5 transformants with 2 wild-type clones of *H.pluvialis* NIES144 as control. Primer pair, tubqf/tubqr, showed there was no detectable genomic DNA contamination in the extracted RNA due to presence of only a single band in amplified products. Because of an intron, genomic DNA contamination would have been indicated by a second band. Primers tubf/aada2r demonstrated transcription of the *aadA* gene driven by the tub promoter in the 5 transformants examined. Spectinomycin resistence in the transformants but not in wild types of *H.pluvialis* was also demonstrated. As shown in Fig 7, growth of the wild type (WT) *H. pluvialis* stopped its growth in liquid media containing 10 ug/ml spectinomycin for 30 days. By contrast, while transforments could grow in media containing 10, 20 or 50 ug/ml spectinomycin. Howevery, cellular growth of the transformants was weakened with elevated antibiotic concentrations, such that in media containing 100ug/ml spectinomycin, their growth was totally inhibited. Through elevated spectinomycin resistance, these results demonstrate the successful expression of the foreign *aadA* gene in the *H.pluvialis* transformants.

**Fig 7.**
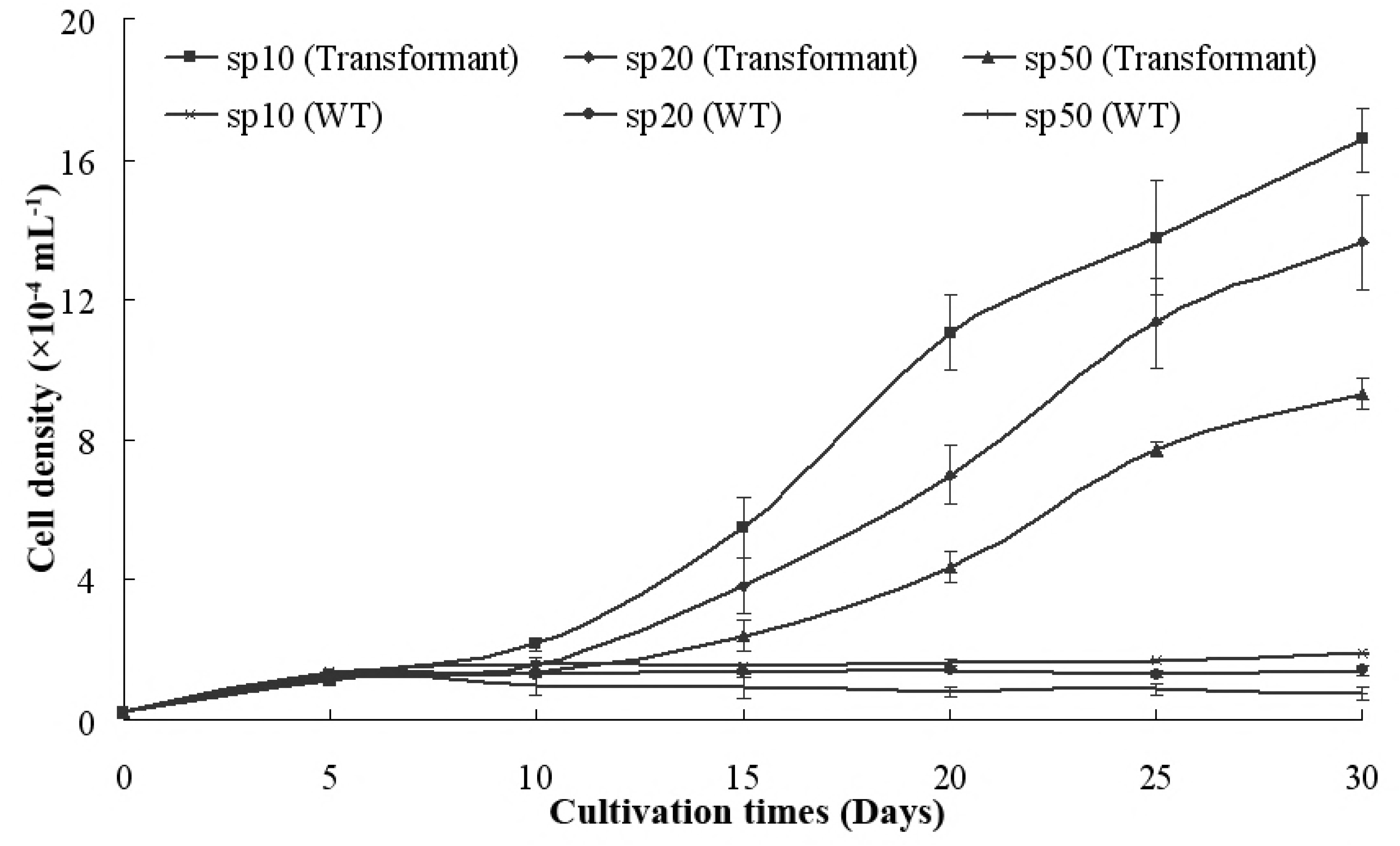
Comparison of antibiotic resistance between transformants and wild types(WT) of *H.pluvialis* NIES144 in suspension cultivations with spectinomycin concentrations of 10 μg/ml (sp10), 20 μg/ml (sp20) and 50 μg/ml (sp50). The data on transformants were from parellel experiments with 3 independent transformants, and 3 parallel wild type samples.

## Discussion

Alpha tubulin (*tub*) is a principal component of microtubules, which are ubiquitous constituents of eukaryotic mitotic spindles, cytoskeleton, cilia and flagella [17]. The coding sequence (CDS) of the *tub* gene in *H.pluvialis* was firstly cloned by Eom et al. in 2006 and its stable expression was shown by transcript analysis [18]. For this current study, the complete sequence of *tub* genes, including exons, introns, 5’ and 3’ flanking sequences were identified for *H.pluvialis* NIES144 and SCCAP K0084. In *H.pluvialis*, the flanking sequences of several genes involved with astaxanthin synthesis, such as bkt (beta-carotene ketolase gene) [19], ipp1(the isopentenyl pyrophosphate isomerase), crtR-B(carotene hydrolase gene) [20] and pds [9], were also cloned and sequenced. But most of the clones only covered the 5’ flanking region. With the development of high throughput transcriptome sequencing, cloning the CDS of functional genes has become a relatively routine task. By contrast, cloning the 5’ and 3’ flanking regions of genes in *H. pluvialis* is still based on genomic walking or screening of genomic libraries. In 2018, the chloroplast genome of *Haematococcus lacustrisis* was deposited in the Genebank using PacBio long-read sequencing technology [21]; however, its nuclear genome is still unreported. This situation, limits the quick access to and application of transcriptional regulatory sequence in the *H.pluvialis* nuclear genome.

An active promoter is the key element in genetic transformation or expression systems. In the tub 5’ flanking region, 1 putative TATA-box, 11 CAAT boxes and some other regulatory transcriptional elements were identified and they were conserved between the 2 *H.pluvialis* strains. Both the TATA box and CAAT box belong to the core promoter elements, responsible for initiation of gene transcription in eukaryote. In the β- carotene ketolase gene promoter of *H.pluvialis* 1 TATA box and 6 CAAT boxes were identified, which showed universal promoter activity and could drive expression of antibiotic resistance genes in *Chlamydomonas reinhardtii* CC-849 [19]. Several putative light-regulated elements were also found in the *tub* promoter, such as ACE motif, CATT motif, GAG motif and T box, which were comparatively less common in the bkt and crtR-B promoters. This might be due to the contitutive-expressing character of the *tub* gene. ABRE and MYB elements, involved in abscisic acid response and drought stress, were also identified in the *tub* promoter, in addition to the bkt [19] or crtR-B [20] promoter.

In eukaryotes, cis-acting elements in 3’ flanking sequences participate in transcription termination, processing and polyadenylation of premessenger RNA [22]. Therefore, the endogenous terminators in 3’ flanking sequences are of equal importance in the construction of transforming vector as the promoter. Knowledge about the poly(A) signals (PAS) is mainly from the research in animals, yeasts and plants [23]. The sequence AAUAAA, the well-known poly(A) signal, was firstly identified in mRNAs of animals. In plants, it was classified as the near-upstream element (NUE) which was an A-rich region of between 6 and 10 nucleotides (nts) [24]. As shown in Fig 2, the typical AATAAA motif existed in the 3’ flanking sequences of *tub* genes of 2 *H.pluvialis* strains. The 3’ flank sequences of the *H. pluvialis tub* genes has a similar architectures to PAS in plants. Consensus signals for plants do include some unusual combinations, such as nebulous UG motifs [25] or the sequence TTGTAA [26] as Far-Upstream Element(FUE), as well as the NUE and a CA sequence at 13 - 30 nts downstream from NUE as the cleavage site(CS) [27]. There was no other information about 3’ flanking region of *H. pluvialis* genes except for the *pds* gene. No similar architecture was found in the *pds* 3’ flanking region: however, this may be due to shortage of sequence information limited to 330 nts limiting the finding of similar elements.

The selectable marker system is key to screening for transformants with stable integration of foreign genes into *H.pluvialis* genome. Transient expression of galactosidase (LacZ) genes respectively driven by a simian virus 40 promoter and by the β-carotene ketolase promoters have been identified; however, no stable transformation was obtained [28]. Only 3 selective toolboxes have shown stable transformation of *H.pluvialis*: *hptII* driven by CaMV 35 promoter in pCAMBIA1302 [11], a mutated endogenous PDS gene in pPlat–pds [9] and pBS–pds [10], and *aadA* driven by an endogenous rbcl promoter in the plamid pHuplus [12]. In current study, 3 antibiotic-resistant genes, *aadA, sh ble* and *hptII*, were each independently fused with the *tub* flanking sequences of *H.pluvialis* NIES144 and were used to transform *H.pluvialis*. Stable transformants with integrated sequences and showing expression of *aadA* gene were obtained; however, constructs tub/hptII and tub/ble failed to produce transformants after multiple attempts. pCAMBIA 1305, has the hptII CDS downstream of the CaMV 35 promoter, was also tested in biolistic transformation of *H.pluvialis*, but it also failed to produce transformants. Similar result was also reported by the reference [10]. The *sh ble* gene has been utilized in the transformation of *Chlamydomonas reinhardii* [29], *Phaeodactylum tricornutum* [15] and *Nannochloropsis oculata* [30]. The *sh ble* cassette driven by the PsaD promoter from *C.reinhardii* was once integrated into the genome of *H.pluvialis*, mediated by pBS–pds. There was confirmed transcriptional expression of the gene but no resistance to zeocin [10]. It was thought that unsuitable codon usage in the gene resulted in insufficient antibiotic-resistance protein expression in transformed cells. Different codon usage may have led to the failed transformation with the tub/ble cassette.

Biolistic transformation is a universal genetic transformation technology [31], which has been used to target to the genomes of both the nucleus [9] or the chloroplasts [12] in *H.pluvialis*. In this paper, the plasmid pEASY-tub/aadA was also transformed into *H.pluvialis* NIES 144 and SCCAP K0084 successfully. M17 tungsten particles were utilized as DNA carriers. An improvement of transformation frequency through the use of gold particles instead of tungsten has been reported by the reference [10]; however it was found that gold particles were easy to adhere to the inner wall of eppendorf tubes leading to some difficulties in the DNA coating process. And transforming efficiencies of 1.2-3.0×10^-5^ per μg DNA of pEASY-tub/aadA were obtained by using cheap tungsten particles, similar with previously reported efficiencies of 1.6×10^-5^ and 2×10^-5^ per μg DNA for the plasmids pHuplus and pBS–pds by golden particles. In a regular procedure, DNA was dissolved at a concentrantion of about 1 μg μL^-1^ for coating, and particles with about 1 μg coated DNA were utilized for 1 bombardment. Multiple bombardments on the same batch of cells were often done to increase the stablility and efficiency of transformation [10]. In this work, increased DNA amount on 1 bombardment also enhanced *H.pluvialis* transformation. The PCR-amplified DNA fragments could also be successfully utilized for biolistic transformation of *H.pluvialis*, though at a lower efficiency than supercoiled plasmid DNA was identified. Using PCR fragments was thought to be timesaving in allowing both functional analysis and construction of modified strains. Similar results with DNA fragments have also been reported in *Phaedactylum tricornutum* [32]. In order to increase the stability of each bombardment, concentrated cells was uniformly pipetted onto a membrane with a diameter of 2.4 cm, which could guarantee all the cells were in the target zone of each bombardment. Moreover, bombarded cells had be washed immediately from the membrane into regeneration medium due to the bad survival of *H.pluvialis* cells on the membrane.

Besides optimizing the bombardment process, it was also important for successful transformation to choose the best culture phase based on the content of flagellate cells in the culture. *H.pluvialis* are differentiated into motile flagellates and non-motile resting cells [33]. Non-motile resting cells had thickened cellulosic cell-walls. When a late-stage culture after 12 days cultivation of *H.pluvialis* SCCAP K0084, containing only about 40% of the cells in the motile stage, was used for bombardment, no transformation was achieved. It was previously that transient expression of lacZ could be observed in motile cells but not in non-motile cells after bombardments [28]. While cell differentiation of *H.pluvialis* could be influenced by many environmental factors, and there are also great differences among different *Haematococcus* species or strains [34]. In this research, enriched flagellates were prepared in BYA medium as previously reported by [16]. The cultures of *H.pluvialis* NIES144 and SCCAP K0084 chosen for stable biolistic transformation both had > 80% of cells as motile flagellate.

Through biolistic transformation, the tub/aadA cassette could be stably transformed into the genome of *H.pluvialis*. Transcripts for the foreign gene and antibiotic resistance were identified in these transformants. The foreign DNA existing as a supercoiled form in the nucleus could be excluded, while no homologous recombination, including single crossover or double crossover, could be detected in transformation with plasmid pEASY-tub/aadA or the tub/aadA PCR fragment. However, there were also no clear data showing existence of homologous recombination in nuclear transformation of *H.pluvialis* according to previous studies [9, 10]. Through homologous recombination, stable disruption can be achieved in targeted genes, and this has been widely utilized in many model species [35]. In eukaryotic microalgae, most transforming DNA integrates at non-homologous or random sites in the nucleus. It was reported that homologous recombination requires a region of homology of no more than 230 bp in *C. reinhardtii* [36]. And there are only 2 reports describing the targeted disruption of genes in the nucleus of *C. reinhardtii* [37] and *Nannochloropsis sp.*[38].

In conclusion, stable nuclear transformation was achieved in *H.pluvialis* NIES144 and SCCAP K0084 through biolistic bombardments, using the *aadA* gene driven by transcriptional elements flanking the endogenous alpha tubulin gene. This approach enhances the tools available to genetically modify *H.pluvialis*. Besides well-established bombarding conditions, 2 cellular factors contributing to the success of *H.pluvialis* transformation were identified in this research, the importance of a high flagellate percentage in the culture and the low survivability of cells on the filter membrane. Establishment of a stable and efficient transforming technology should facilitate researches in metabolic engineering and synthetic biology in *H.pluvialis*.

## Acknowledgements

The project was supported by 2016 Sino-Thai Cooperation Project by National Natural Science Foundation of China (Grant No. 51561145015) and financial support from Shandong Provincial Key Laboratory of Energy Genetics.

